# A mouse model of oral contraceptive exposure: depression, motivation, and the stress response

**DOI:** 10.1101/2023.01.27.525969

**Authors:** Kristen M. Schuh, Jabir Ahmed, Esther Kwak, Cecilia X. Xu, Tronjay T. Davis, Chloe B. Aronoff, Natalie C. Tronson

## Abstract

Hormonal contraceptives, including oral contraceptives (OCs), regulate hormonal cycles and broadly affect physiological processes, including stress responsivity. Whereas many users describe overall improved mood, up to 10% of OC users experience adverse effects, including depression and anxiety. Given the link between regulation of hypothalamic-pituitary-adrenal (HPA) axis, stress exposure, and risk for depression, it is likely that OC-effects on stress mediate increased risk or increased resilience to these disorders. In this study, we developed and characterized a tractable mouse model of OC exposure with which to identify the mechanisms underlying OC modulation of brain, behavior, and mood. Specifically, we aimed to determine whether translationally relevant doses of OC-hormones in mice mimic changes in stress responsivity observed in humans taking OCs and describe behavioral changes during OC exposure. Young adult female C57Bl/6N mice received daily ethinyl estradiol (EE) and levonorgestrel (LVNG) in 10% sucrose, EE and drospirenone (DRSP) in 10% sucrose, or 10% sucrose alone. Translationally relevant doses of EE+LVNG-exposure, but not EE+DRSP, suppressed the acute stress response, consistent with effects observed in human OC users. EE+LVNG caused a specific anhedonia-like effect, without broad changes in stress-coping behavior, other depression-like behaviors, or anxiety-like behaviors. The suppression of regular estrous cycling, together with the blunting of the corticosterone response to acute stress, demonstrate the utility of this model for future studies to identify the mechanisms underlying OC interactions with stress, motivation, and risk for depression.

## INTRODUCTION

Hormonal contraceptives, including oral contraceptives (OCs, “the pill”), IUDs (e.g., Mirena), implants (e.g., Nexplanon), and the vaginal ring, are a critical part of healthcare, providing an unprecedented level of reproductive control and, in turn, economic and educational benefits for families. In addition, because OCs blunt ovarian cycles, they also exert health benefits, including alleviation of menstrual symptoms and treating premenstrual dysphoric disorder, controlling acne, and reducing the risk of ovarian, epithelial, and endometrial cancers (Havrilesky et al., 2013; Michels et al., 2018; Murphy et al., 2017; Schindler, 2013). Despite the myriad of benefits for many users, hormonal contraceptives have some side effects, and adverse effects on mood, remains the most common reason for cessation of use (Lewis et al., 2019; Sanders et al., 2001). At least 4-10% of users – or up to 30 million individuals at any given time – experience increased risk for mood disorders (Poromaa & Segebladh, 2012).

Whereas OCs increase depression and anxiety for some users, others experience beneficial effects on mood (Keyes et al., 2013; Porcu et al., 2019; Poromaa & Segebladh, 2012; Tronson & Schuh, 2022). These conflicting findings are likely due to individual differences in mood-related risk factors, differences between OC formulations, and different inferences from analyses that focus on group averages versus those that assess individual risks or changes across time (Beltz, 2022). This raises the question of how to predict who is likely to experience beneficial effects and who is at risk for detrimental effects – but to do that we need to better understand how OCs affect the brain. Here, we begin to characterize a new mouse model with which to identify mechanisms by which OCs contribute to regulation of stress and risks for anxiety and depression.

Developing animal models is a necessary step towards understanding how OCs modulate the brain and behavior (Hilz, 2022; Tronson & Schuh, 2022). Previous studies in rodents have successfully modeled OC exposure in rodents using daily hormone injections, implants, or daily oral gavage protocols to model suppression of ovulation, extensions of fertility, modulation of social and affective behaviors, cognition, and changes in neurochemistry (Allaway et al., 2021; Carrington & Bailey, 1985; Isono et al., 2018; Lacasse et al., 2022; Porcu et al., 2012; Simone et al., 2015). Here, we build on these studies to design a mouse model specifically targeted to mimic OC exposure, their effects on the hypothalamic-pituitary-adrenal (HPA) axis, and subsequent changes in risk / resilience to depression- and anxiety-like processes.

Across studies and across individual OC users, one reliable physiological effect is the suppression of the acute stress response (Kirschbaum et al., 1995; Mordecai et al., 2017; Tronson & Schuh, 2022). Given that stress and dysregulated stress-related signaling play a key role in depression and anxiety, changes in the HPA axis as a consequence of OC use is particularly interesting for identifying how OCs influence risk and resilience to depression (Slattery & Cryan, 2017; Tafet & Nemeroff, 2016).

The main goal of this study was to determine whether translationally relevant doses of ethinyl estradiol + levonorgestrel (EE+LVNG) in mice mimic the suppression of HPA-axis response to acute stress observed in humans, and thereby provide a basis for future mechanistic studies on the interactions of OCs, stress, and depression. Here, we assessed stress-coping, anhedonia-, anxiety-, and other depression-like behaviors. These mouse behaviors are sensitive to stress and dysregulation of HPA-axis and have been used extensively to study behavioral, circuit, neurochemical, and molecular changes associated with stress- and depression-related phenotypes (Bangasser & Valentino, 2014; Commons et al., 2017; Kolber et al., 2008; McEwen, 2000; Serchov et al., 2016; Tafet & Nemeroff, 2016).

Based on previously observed effects in humans (Hertel et al., 2017; Kirschbaum et al., 1995; Merz et al., 2012) and rats (Allaway et al., 2021; Carrington & Bailey, 1985; Follesa et al., 2002; Lacasse et al., 2022; Porcu et al., 2012; Simone et al., 2015), we hypothesized that a translationally relevant dose of orally administered contraceptive hormones will suppress acute stress responses and trigger increases in anxiety- and, depression-like behaviors. Our main findings supported the suppression of corticosterone in response to stress, and showed a specific anhedonia-like effect, without broad depression-like or anxiety-like effects.

## METHODS

### Animals

94 adult female C57Bl6/N mice (9-10 weeks on arrival) from Envigo (Indianapolis, IN), used across four cohorts, Mice were individually housed in the colony room with access to standard chow and water *ad libitum*. Mice were allowed at least 7 days to acclimate to the colony room prior to behavioral experiments or hormone exposure. The colony room, adjacent to behavioral testing rooms, was maintained at 20 ± 2 °C with a 12h 0700: 1900 light/dark cycle (lights on at 0700 h). All protocols and treatments were approved by the University of Michigan Institutional Animal Care and Use Committee.

### Hormone and control treatments

were given for at least 2 weeks prior to behavioral testing and throughout all behavioral and stress testing and tissue collection. Control mice received 250μL of 10% sucrose. Hormone treated groups received 250μL of 10% sucrose with a suspended combination of ethinyl estradiol (EE, 0.01875μg/20g mouse; approximately 0.9375μg/kg) and progestin—either the 2nd generation levonorgestrel (LVNG, 0.75μg/20g mouse, or 37.5μg/kg) or the 4^th^ generation progestin drospirenone (DRSP, 3.75μg/20g mouse, or 187μg/kg). These doses were based on prior publications demonstrating suppression of ovulation in mice (Isono et al., 2018); and calculated to approximate human doses, with corrections for body surface area and metabolism (Nair & Jacob, 2016).

For all groups, 250μL was delivered daily in small cell culture dishes in the animals’ home cage prior to lights out (1700-1900h). Mice were observed, and the solution was consumed within minutes.

### Experimental Approach (Figure 1)

In all cohorts, animals received hormone/control treatments each day for the duration of the experiment. Behavioral tasks began at least 2 weeks after the start of hormone/control treatments; and all tests were conducted in the light cycle, at least 16 hours after hormone treatment to avoid acute effects of hormone (or sucrose) ingestion. Behavioral tasks were conducted in order from least stressful to most stressful to minimize stress effects on future behavioral tasks. The data from this experiment comes from 4 different cohorts of animals. In cohort 1, we assessed suppression of estrous cycle and acute stress response; cohort 2, mice were tested on open field test, sucrose preference, and forced swim test; cohort 3, mice were tested on taste preference, sucrose preference, instrumental training, and progressive ratio; and cohort 4, mice were tested on novelty suppressed feeding, light-dark emergence, and fear conditioning.

**Figure 1.**
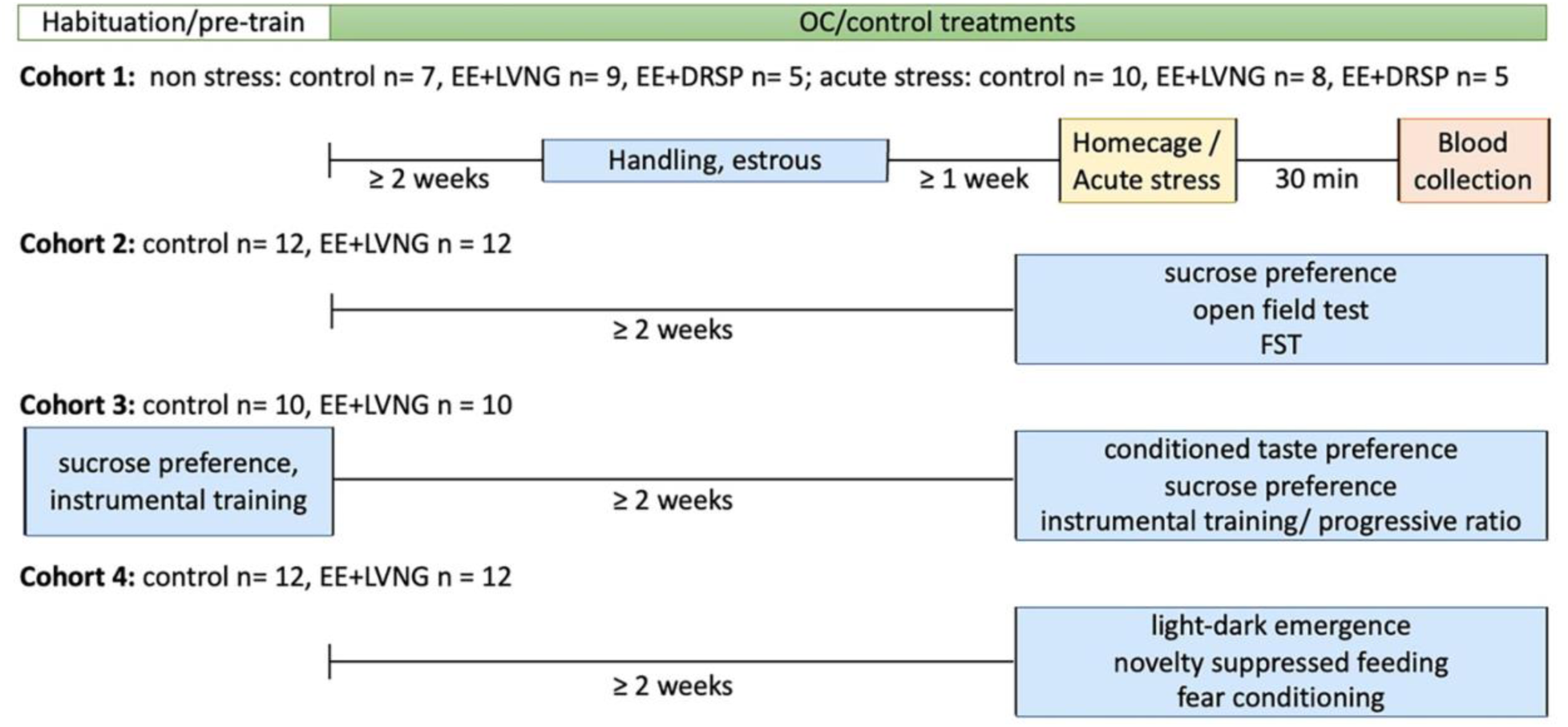
Timeline of experiments. Four separate cohorts of mice were used throughout this project.

### Estrous Cycle

was assessed using vaginal cytology and visual assessments (Caligioni, 2009; Keiser et al., 2017) in cohorts 1 and 2. Unpaired t-tests were used to compare the number of estrous phase shifts between control and EE+LVNG treatment groups. In order to reduce potential stress confounds, we did not assess estrous cycles in cohorts 3 or 4.

### Weights

were measured weekly throughout the experiment in cohort 2. A repeated measures ANOVA was used to compare weights of control (n=12) vs EE+LVNG-treated (n=12) animals across days.

### Acute Restraint stress

Animals were briefly anesthetized using 0.1 mL isoflurane in a drop box and placed prone on a flat surface. Tape was applied to the elbow joints of the forelimbs and the knee joints of the hind limbs so that animals were restrained for 1 hour (Tronson et al., 2010). After which, the tape was carefully removed to minimize unnecessary discomfort, and animals were returned to their home cage.

### Blood collection

Thirty minutes after the end of the acute stressor, mice were deeply anesthetized with an intraperitoneal injection of 250 mg/kg of Avertin (2,2,2-tribromoethanol), and blood was collected from the heart using cardiac puncture. Blood was centrifuged at 15294 g (12000 RPM) for 15 minutes. Serum was removed and stored at −20°C.

### Corticosterone ELISA

(Cat # K014-H1, Arbor Assays, Ann Arbor, MI) was used according to the manufacturer’s instructions (Epoch plate reader, Biotek). A 2-way ANOVA was conducted with treatment and stress exposure as factors. Post-hoc tests with Bonferroni correction were conducted to determine the effects of stress within treatment groups.

### Selected Behavioral tasks

were used to assess depression and anxiety endophenotypes. We selected well-established behavioral tasks to assess specific processes involved in depression and anxiety disorders. For depression-like behaviors we used sucrose preference to assess anhedonic processes, forced swim test to assess stress-coping strategies, and novelty suppressed feeding to assess motivational conflict (Commons et al., 2017; Dulawa & Hen, 2005; Liu et al., 2018). For anxiety-like behaviors we used open field test to assess locomotor activity, exploration, and willingness to explore an open area (Seibenhener & Wooten, 2015), light-dark emergence to examine risk-assessment behaviors (Arrant et al., 2013), and context fear conditioning to assess acquisition of a new aversive memory and fear-related behaviors (Kim & Cho, 2020). Additional tasks (e.g., conditioned taste aversion, instrumental learning, and progressive ration tests) were used to examine alternative explanations for observed behavioral changes (Kislal & Blizard, 2016; Sharma et al., 2012).

#### Apparatus

Behavioral boxes (8.5”x 7.12” x 5”) fitted with lixit bottles, within sound attenuating cubicles (MedAssociates, 25” x 17” x 17.5”) were used for the sucrose preference and conditioned taste avoidance tests. These MedAssociates boxes fitted with nose-poke holes and pellet dispensers were used for instrumental training, and progressive ratio tests. Contextual fear conditioning was conducted in rectangular Perspex conditioning boxes (9 3/4′′ × 12 3/4′′ × 9 3/4′′; MedAssociates, VT) with evenly spaced grid floors within sound-attenuating chambers, and NIR VideoFreeze (MedAssociates, VT) was used to assess freezing. A square open field (1m x 1m) was used for the open field, novelty suppressed feeding, and light/dark emergence tests. These tests were recorded and automatically scored using Ethovision XT (v.16; Noldus Information Technology, VA).

#### Sucrose preference

Mice were habituated to 2% sucrose in their home cage and in the MedAssociates boxes to prevent neophobia of the sucrose or context from suppressing drinking (Strekalova & Steinbusch, 2010). Each box was equipped with two Lixit bottles containing 1-2% sucrose. To avoid acute effects of hormone, all habituation and tests were in the morning (0900-1100h) after overnight water fasts. At test, mice were placed into the identical boxes with a 2-bottle choice between 1-2% sucrose and water for 30-min sessions. A second day of testing counterbalanced for side biases. In cohort 2, mice were tested on preference for 2% sucrose. In cohort 3, mice were first matched for sucrose preference prior to OC treatment and tested with 1% sucrose. In both cohorts, the number of licks were automatically detected using MedPC software (MedAssociates, VT), and preference was calculated as a percentage of sucrose *vs* total licks [100*# sucrose licks/(#sucrose + #water licks)]. Sucrose preference was assessed using unpaired one-sample t-tests to compare the preference ratios of each treatment group to a theoretical mean of 50. Mice that had fewer than 10 total licks were excluded from the sucrose preference analysis. A 30-min sucrose preference test was chosen instead of an overnight intake so that we were avoiding the acute changes in the hours immediately after EE+LVNG treatment, and any acute effects of 10% sucrose ingestion.

#### Conditioned Taste Aversion Task

was used to assess whether EE+LVNG administration caused a taste aversion to sucrose (Kislal & Blizard, 2016) as a possible explanation for disrupted sucrose preference in EE+LVNG-treated mice. On the first 2 consecutive days of hormone exposure, all mice received 10% sucrose paired with Flavor A at the beginning of the light cycle (0900-1100h) and 10% sucrose with EE+LVNG (controls received 10% sucrose alone) combined with Flavor B at the end of the light cycle (1700-1900h). Flavors were orange and cucumber, diluted at 1:1000 (SodaStream), and counterbalanced across animals. A two-bottle choice was used (1600-1800h, cucumber vs orange, 1:1000 dilution in tap water), and the number of licks on each bottle was measured. Preference ratio was calculated as described above. Unpaired one-sample t-tests were conducted for each treatment group to compare the group mean preference ratio to a theoretical mean of 50.

#### Novelty Suppressed Feeding

was used to assess the conflict between motivation to eat and avoidance of novelty (Dulawa & Hen, 2005). After an overnight fast (testing occurred between 1000-1400h), mice were placed into the open field apparatus with a pellet of standard chow placed on a circular piece of filter paper (4.5 in diameter) in the center. Latency to step on the filter paper and explore the food was automatically recorded. Differences in latency to the food pellet between treatment groups were assessed using an unpaired one-sample t-test.

#### Forced swim test

is a measure of active *vs* passive stress-coping behaviors, which is sensitive to changes in HPA-axis functioning (Commons et al., 2017). Mice were placed in a 2000 mL beaker filled 2/3 with room temperature water (22° C +/- 1°C) for 6 minutes (testing occurred between 1000-1500h). After the test, animals were dried off with paper towels and placed in their home cage with a heating pad for 10 minutes (Sunbeam, 43.3°C). Time spent immobile (no movement other than that required to stay afloat) was hand scored during minutes 2-6 of the task by three researchers masked to condition. An unpaired t-test was conducted to compare time spent immobile between EE+LVNG and control treatment groups.

#### Light/Dark Emergence

was used as a sensitive measure of anxiety and risk assessment (Arrant et al., 2013). Testing was conducted between 1000-1600h. Mice were placed in a black box (2.5 x 5 x 3 in) in the center of the open field. The door was opened, and latency for nose, forepaw, and full body to emerge from the box was hand scored by two researchers who were blind to experimental conditions. Unpaired t-tests were used to compare first nose emergence, first forepaw emergence, and whole-body emergence between treatment groups.

#### Open field test

was used to assess exploratory and anxiety-like behavior (Seibenhener & Wooten, 2015). Animals were placed in an open field for 10 minutes (testing occurred between 1000-1500h). Total distance travelled and cumulative duration at the field edges and center were measured. Less time in the center of the arena is an index of increased anxiety-like behavior. Unpaired t-tests were used to compare distance travelled and time-in-center between treatment groups.

#### Context Fear Conditionin

was used to assess fear learning (Kim & Cho, 2020). The task was conducted between 1200-1600h as previously described (Keiser et al., 2017). Mice were placed into the chamber for 3 minutes, followed by a 2s, 0.8mA footshock.

Mice were returned to their home cage in the colony room immediately following training. 24-hours later, animals were placed into the identical chamber for 3-minutes. Locomotor activity and freezing were automatically assessed throughout training and testing via VideoFreeze (MedAssociates, VT). The chambers were cleaned with 70% ethanol between each mouse. Unpaired t-tests were used to compare locomotor activity during training, shock reactivity during training, and freezing during test between control vs EE+LVNG-treated groups.

#### Instrumental Training and Progressive Ratio

were used to assess appetitive learning and motivation (Sharma et al., 2012). To avoid any group differences in food intake during initiation of training, mice were pre-trained prior to OC exposure and matched EE+LVNG vs control groups on performance after day 4. Mice were food restricted up to 5 days/week with *ad lib* food at least 2 days a week to maintain at 85% of their free-feeding weight. All testing occurred between 0900-1200h. Mice were habituated to grain food pellets (F0163, Bio-Serv) in a dish in their home cage and subsequently trained to nosepoke for the food pellet reward. Animals were trained on an FR1 schedule to a criterion of at least 20 total food pellets and 75% accuracy for three consecutive days, followed by an FR4 schedule to the same criterion. Mice then underwent a progressive ratio task using a grain pellet reward, in which the number of nosepokes required to earn one food pellet increased exponentially ([5*e*^(*reward number* ×0.2)^] − 5) (Richardson & Roberts, 1996). Mice were retrained and tested according to the previous schedules with chocolate-flavored sucrose pellets (F05301, Bio-Serv). An unpaired t-test was conducted to compare the number of days it took for the animals in each treatment group to reach criteria. A Kaplan-Meier survival analysis was used to compare the number of control vs EE+LVNG-treated animals that reached criteria across days. Unpaired t-tests were conducted to compare break points across treatment groups in the progressive ratio task with the grain pellets and the progressive ratio task with the chocolate pellets.

## RESULTS

### Daily EE+LVNG exposure disrupted the estrous cycle in mice

Vaginal cytology was conducted, and cell types were imaged to determine phase of the estrous cycle (as previously described in Caligioni, 2009; Keiser et al., 2017). Figure 2A displays representative images of cell types from each phase. Phases were graphed and control animals were compared to EE+LVNG-treated animals (Figure 2B). EE+LVNG-treated animals experienced fewer estrous phase shifts (t_14_ = 6.75, p < 0.0001, η_p^2^_ = 0.77; Figure 2C); therefore, EE+LVNG exposure disrupted the estrous cycle. In cohort 2, neither control nor EE+LVNG-treated mice showed reliable cycling.

**Figure 2.**
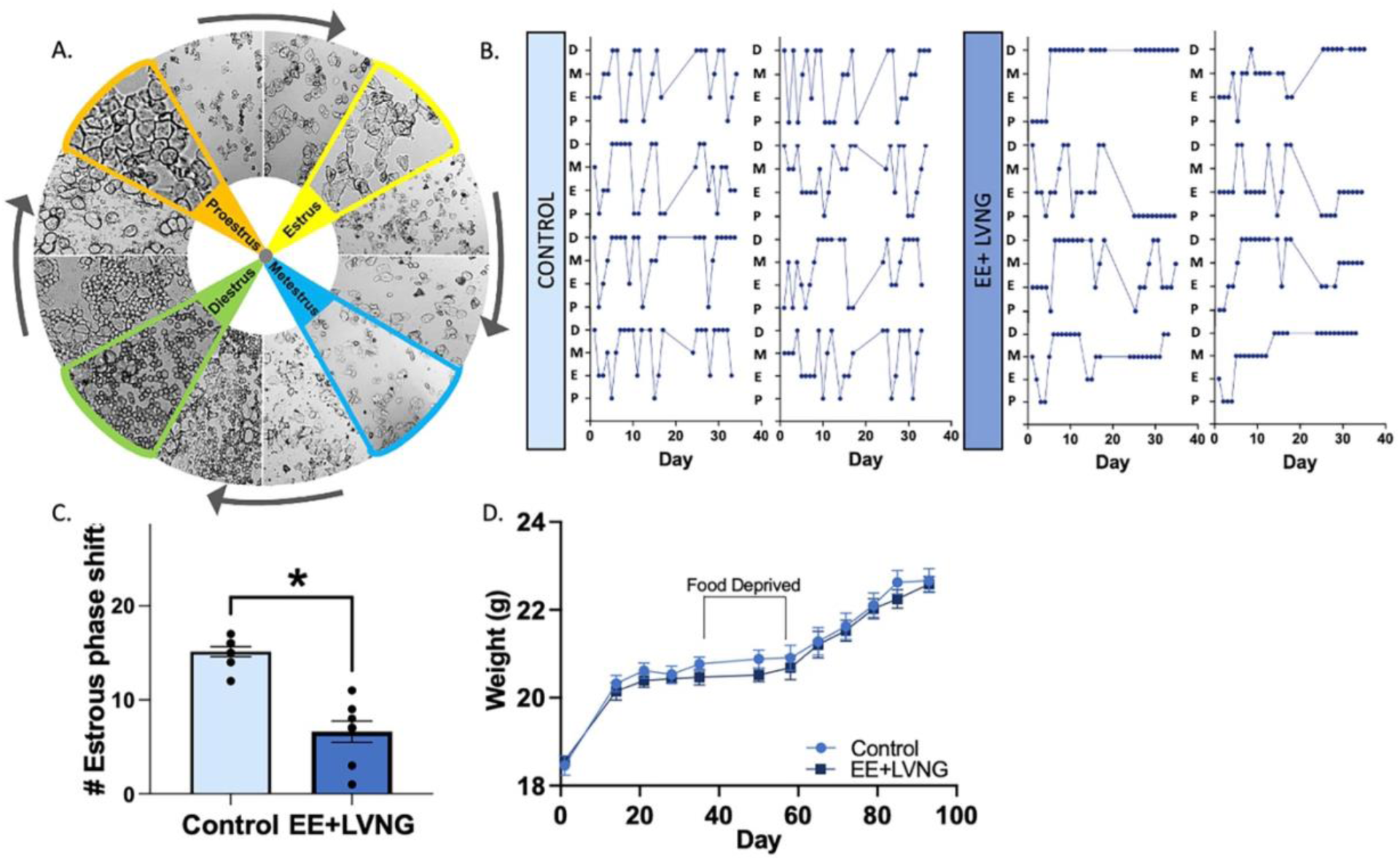
Estrous cycle and weight tracking. (A) Representative images of vaginal smear cell types as they cycle clockwise through each phase of the estrous cycle. P: proestrus, mostly nucleated epithelial cells. E: estrus, mostly cornified cells. D: diestrus, mostly leukocytic cells. M: metestrus, a mix of all three cell types. (B) Visualization of how control (n=8) and EE+LVNG (n=8) mice from cohort 1 cycled through the four phases of the estrous cycle. (C) Control mice in cohort 1 seemed to cycle through all four phases, while EE+LVNG-treated mice did not, * p < 0.0001. (D) Average animal weights throughout the experiment from cohort 2 (control n= 12, EE+LVNG n= 12). The bar represents when the animals were food deprived due to behavioral training. Both treatment groups gained weight during free-feeding and maintained weight during food restriction.

### EE+LVNG exposure did not cause weight gain

The weights of EE+LVNG-treated animals in cohort 2 did not differ from control animals throughout the experiment (Figure 2D). Both groups gained weight during free-feeding and maintained weight during food restriction. We observed a main effect of time (F_11,231_ = 107.1, p < 0.0001, η_p^2^_ = 0.84), but no effect of treatment and no time x treatment interaction (treatment: F_1,21_ = 0.46, p = 0.50; interaction: F_11,231_ = 0.55, p = 0.87). Thus, all animals gained weight across the experimental timeline with no difference as a consequence of EE+LVNG exposure.

### EE+LVNG blunted the acute stress response

EE+LVNG resulted in a significant blunting of the corticosterone response to an acute stressor (Main effect Stress: F_1,38_ = 11.568, p =0.002, η_p^2^_ =0.23; Hormone: F_2,38_ < 1; Stress x Hormone: F_2,38_ = 3.96, p = 0.027, η_p^2^_=0.17; Figure 3). Whereas control mice and EE+DRSP-treated mice, an additional control showed a robust increase in circulating corticosterone 30-min after stress (p = 0.02), EE+LVNG-exposed mice failed to do so (p = 0.88). There were no significant differences between the OC-treated groups and control mice in corticosterone levels of non-stressed mice (Control *vs* EE+LVNG: p = 0.44; Control *vs* EE+DRSP: p = 1.00; EE+LVNG *vs* EE+DRSP p = 0.45).

**Figure 3.**
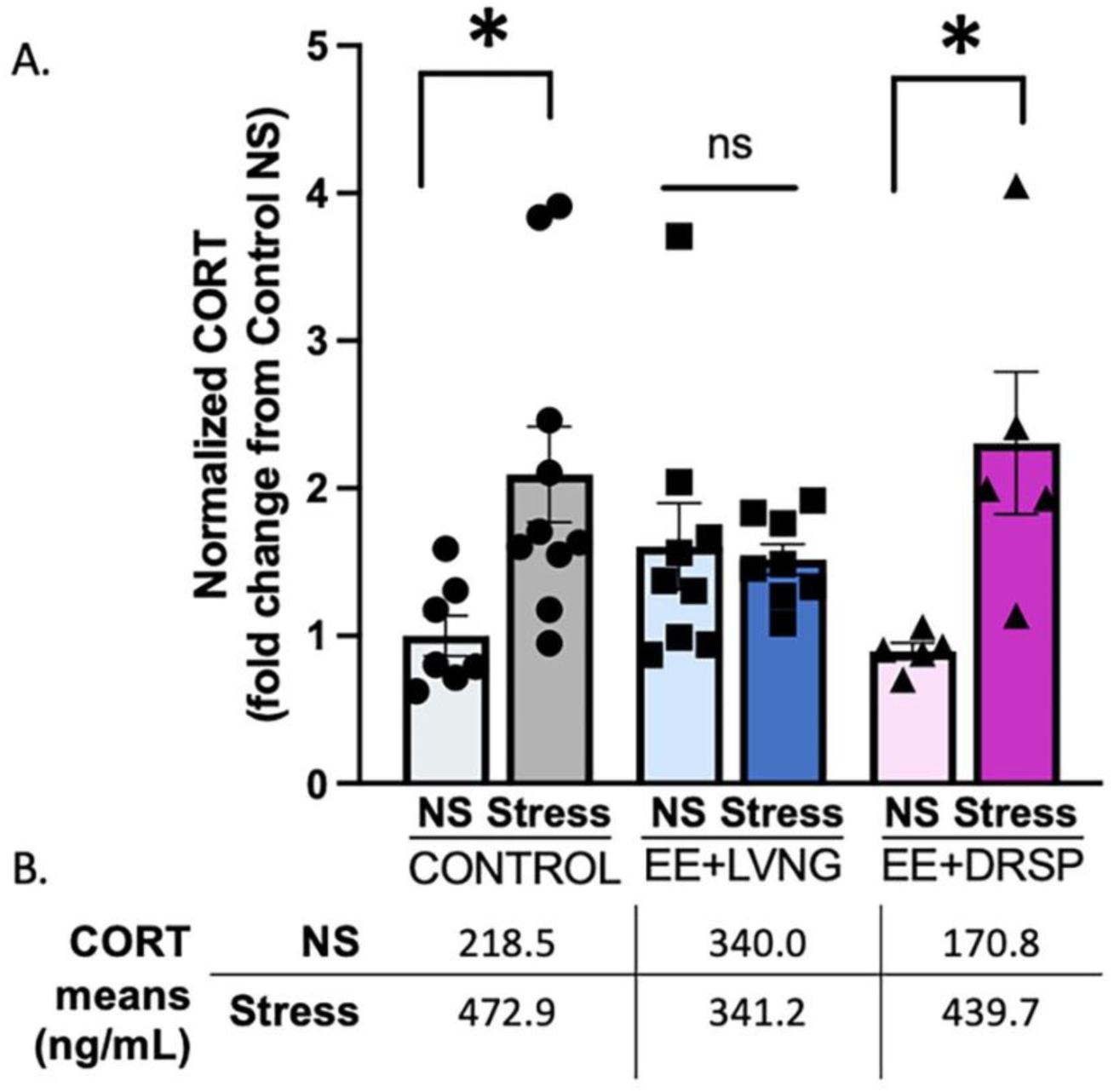
EE+LVNG blunts the acute stress response. (A) Normalized corticosterone levels (fold change from Control NS) after no stress (ns) or acute restraint stress (stress). EE+ LVNG treatment resulted in a blunting of the corticosterone response. However, control and EE+DRSP mice showed a robust corticosterone response to acute stress. *p<0.05. control ns n = 7, control stress n= 10, EE+LVNG ns n =9, EE+LVNG stress n= 8, EE+DRSP ns n= 5, EE+DRSP n= 5. (B) Corticosterone (ng/mL) means.

### EE+LVNG exposure decreased sucrose preference

We assessed preference for sucrose in EE+LVNG and control animals. In two independent cohorts, Control animals, as expected, showed a robust preference for sucrose (cohort 2: t_8_ = 4.871, p = 0.001, η_p^2^_ = 0.75; cohort 3: t_9_ = 7.86, p < 0.0001, η_p^2^_ = 0.87; Figure 4A, 4B); whereas, EE+LVNG-treatment animals showed no preference (cohort 2: t_5_ = 0.08, p = 0.94; cohort 3: t_9_ = 1.77, p = 0.11). Overall consumption during the sucrose preference did not significantly differ between EE+LVNG-treated and control animals (cohort 2: t_13_ = 1.51, p = 0.16, cohort 3: t_18_ = 0.79, p = 0.44; Figure 4A, 4B). When comparing sucrose preference data between the two cohorts, there was no effect of cohort (F_1,31_ = 0.27, p = 0.61) or cohort x OC interaction (F_1,31_ = 1.03, p = 0.32); however, there was a main effect of OC (F_1,31_ = 7.44, p = 0.01, η_p^2^_ = 0.19), demonstrating a decreased preference for sucrose in EE+LVNG-treated mice. In cohort 2, 9 out of the 24 animals did not reach the 10-lick minimum (3 control and 6 EE+LVNG-treated animals) and were therefore excluded in the analyses. In cohort 3, all 20 animals reached the 10-lick minimum. Due to results from both of these distinct cohorts, we show that we can replicate these findings in different groups of animals.

**Figure 4.**
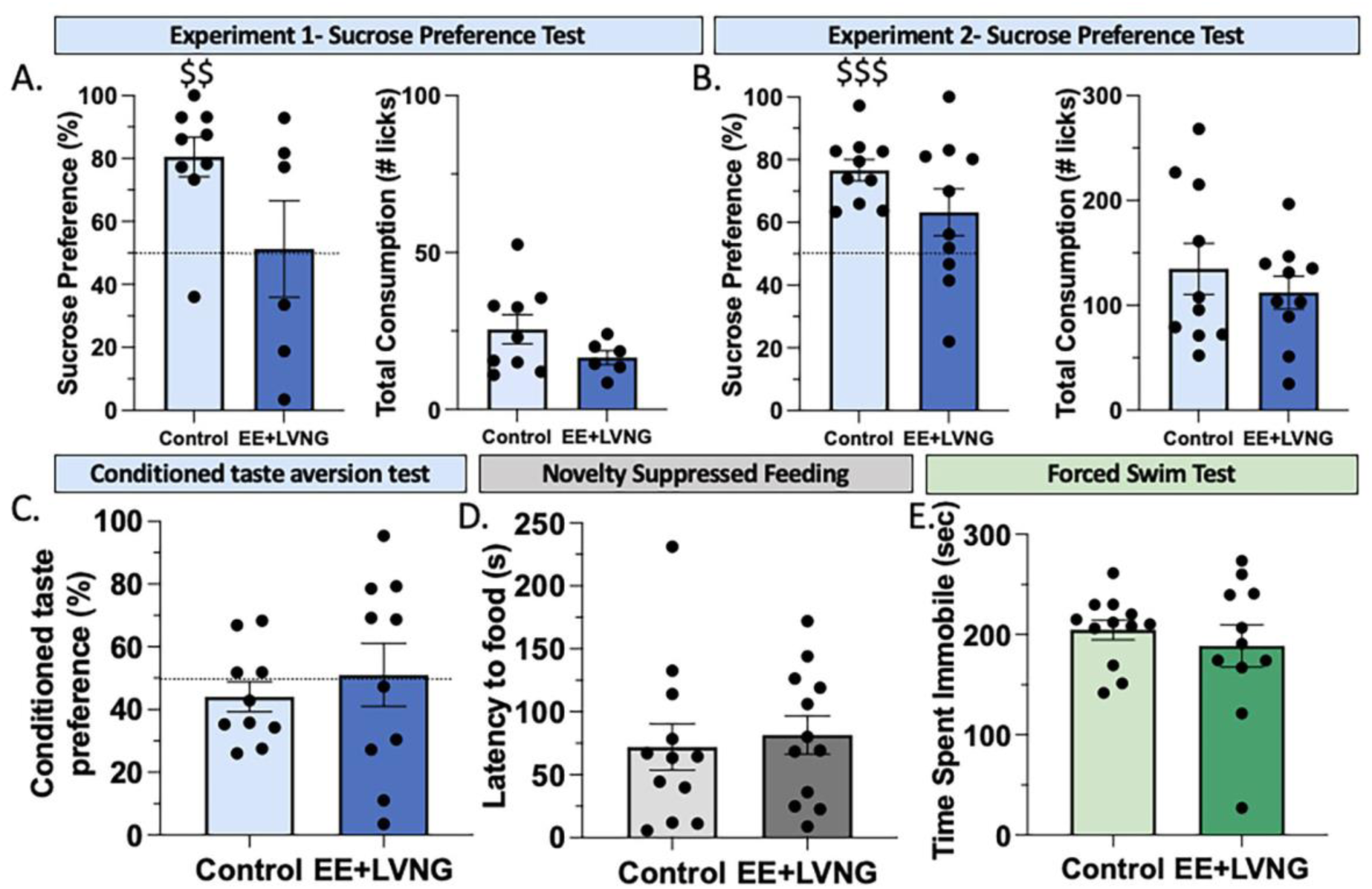
Behavioral tests of depression-like endophenotypes and stress-coping behaviors. (A, B) EE+LVNG mice in two independent cohorts (cohort 2: control n=9, EE+LVNG n=6; cohort 3: control n=10, EE+LVNG n=10) of animals showed a disruption of preference for 1-2% sucrose compared to tap water. $$ p < 0.001 *compared to a theoretical mean of (cf)* 50% $$$ p < 0.0001 *cf* 50%. There was no difference between groups in overall consumption in the sucrose preference test in either cohort. (C) There was no evidence of a taste aversion to the daily oral EE+LVNG treatment (cohort 3: control n=10, EE+LVNG n=10). (D) There was no difference between control and EE+LVNG in latency to approach a food pellet in the novelty suppressed feeding task (cohort 4: control n=12, EE+LVNG n= 12), nor (E) time spent immobile in the forced swim test (cohort 2: control n=12, EE+LVNG n= 11). Light bars: Control-treated mice, Dark bars: EE+LVNG-treated mice.

### EE+LVNG does not cause conditioned taste aversion to sucrose

To determine whether the decreased sucrose preference in EE+LVNG-treated mice might be due to aversion caused by hormone-delivery in sucrose, we conducted a conditioned taste aversion task. There was no evidence of a taste aversion for control (t_9_ = 1.24, p = 0.25) or EE+LVNG-treated animals (t_9_ = 0.11, p = 0.92; Figure 4C), and EE+LVNG mice did not differ from control animals (t_18_ = 0.6313, p = 0.54). Because consumption of EE+LVNG in 10% sucrose did not cause a taste aversion, this is unlikely to be a contributor to decreased sucrose preference in EE+LVNG mice.

### EE+LVNG exposure did not cause stress-coping or anxiety-like behaviors

In contrast to the anhedonia-like effect in EE+LVNG-treated mice, we did not observe evidence of EE+LVNG-related differences in the novelty suppressed feeding task, where latency to approach food did not differ between groups (t_22_ = 0.40, p = 0.69; Figure 4D), nor the forced swim test, where time spent immobile was similar between control and EE+LVNG-treated mice (t_14.2_ = 0.68, p = 0.50; Figure 4E). Together, these data demonstrate that EE+LVNG causes specific anhedonia-like changes. EE+LVNG exposure did not affect exploratory or anxiety-like behavior in the open field task, as neither distance traveled (t_21_ = 0.96, p = 0.35; Figure 5A) nor time spent in the center of the open field (t_21_ = 0.04, p = 0.97; Figure 5B), differed between treatment groups. In the light-dark emergence task, EE+LVNG-exposed animals had a longer latency for forepaw emergence (t_17_ = 2.09, p = 0.05, *η*_p2_= 0.21), but showed no difference in first nose emergence, t_22_ = 1.17, p = 0.25, or whole-body emergence, t_22_ = 0.99, p = 0.33 (Figure 5C, 5D, 5E). This suggests that EE+LVNG does not increase anxiety; however, there might be subtle changes in exploratory or risk assessment behaviors which will be an important avenue for future work.

**Figure 5.**
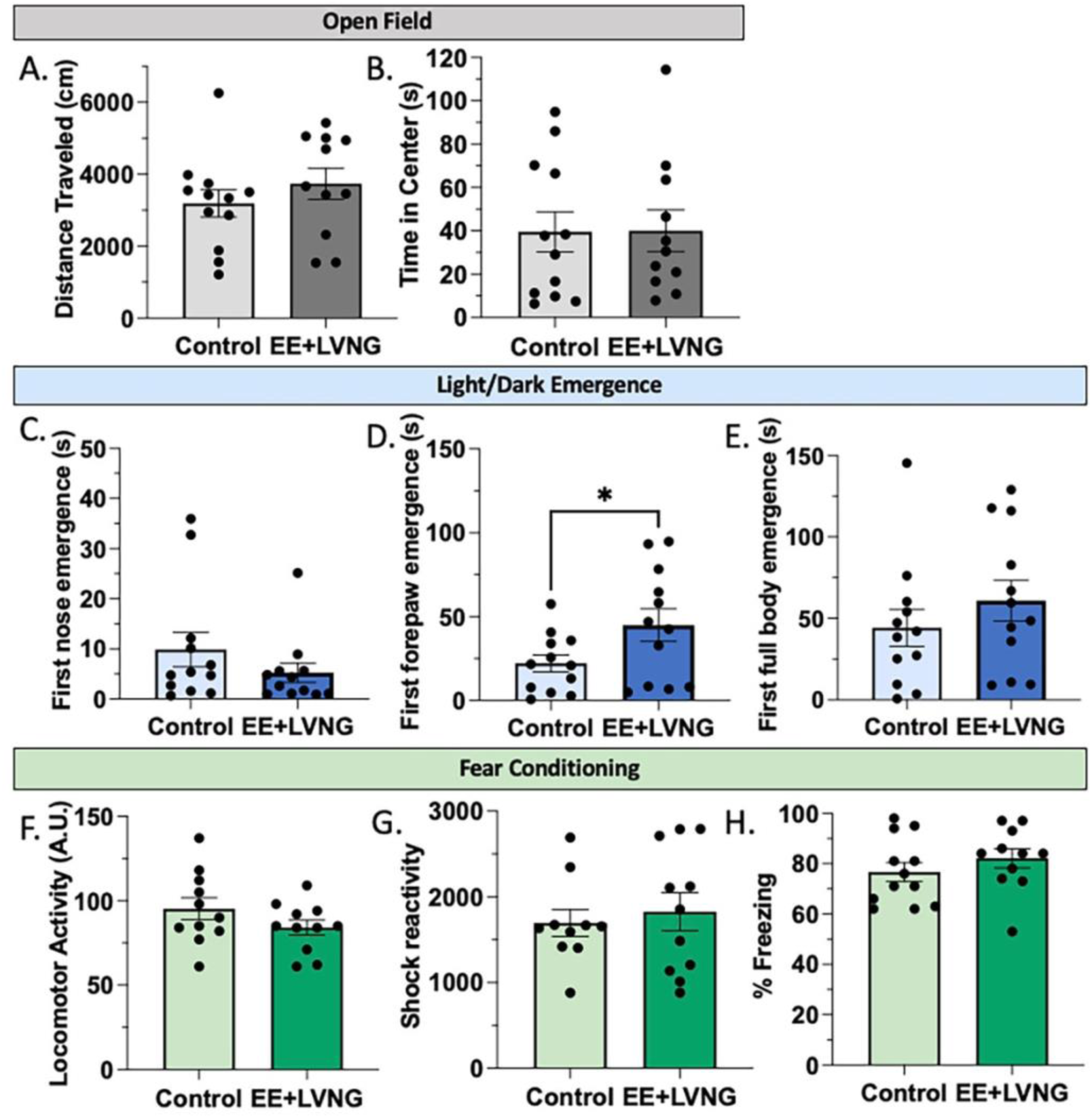
EE+LVNG exposure caused mild changes in risk assessment behavior, without overall anxiety-like effects. (A, B) Neither distance traveled, nor time spent in the center of the open field differed between treatment groups in the open field task (cohort 2: control n=12, EE+LVNG n= 11). (C) EE+LVNG-treated (n=12) and control (n=12) mice had no difference in latency for first nose emergence during the light/dark emergence task; however, (D) EE+LVNG treated mice had a longer latency for first forepaw emergence. *p <0.05 (E) There was no difference in latency of first full body emergence. EE+ LVNG exposure had no effect on (F) locomotor activity during training, (G) shock reactivity, nor (H) percent freezing during the context test (cohort 4: control n=12, EE+LVNG n=12). Light bars: Control-treated mice, Dark bars: EE+LVNG-treated mice.

To assess hippocampal-dependent memory for stress-related information, we used context fear conditioning. There was no effect of EE+LVNG exposure on locomotor activity during training (t_20_ = 1.44, p = 0.17; Figure 5F), animals exhibited no substantial freezing to the context during training, and there was not a difference in shock reactivity, (t_19_ = 0.47, p = 0.64; Figure 5G). There were no differences in freezing (t_21_ = 1.00, p = 0.33) during the context test (Figure 5H). Thus EE+LVNG did not disrupt locomotor activity, response to a noxious stimulus, context fear conditioning, or freezing behavior. Together, these results demonstrate the specificity of EE+LVNG exposure for anhedonia-like processes.

### EE+LVNG did not disrupt instrumental learning or progressive ratio responding

We trained mice on a food-reinforced instrumental task, followed by a progressive ratio test to determine whether EE+LVNG decreased instrumental learning or motivation. All mice acquired the instrumental task, and EE+LVNG-treated animals did not differ from controls in the number of days to reach criteria. Kaplan-Meier survival test demonstrated that survival curves for Control versus EE+LVNG-treated groups did not differ (χ^2^_1_ = 0.009, p = 0.924; Figure 6A). We assessed the number of active and inactive nosepokes during the final three days of instrumental training prior to the progressive ratio. Over the last three days of training with grain pellets, where all animals were performing to criterion, EE+LVNG-treated mice showed a small but significantly higher number of active nosepokes (Treatment: F_1,18_ = 5.40, p = 0.032, η_p^2^_ = 0.231) but no significant effect across day (Day x Treatment, F_2,36_ = 3.00, p = 0.093; Day: F_2,32_ = 3.058, p = 0.088). This effect was driven by the 3^rd^-to-last day of training, with no differences between groups on the last or second to last days (Control vs EE+LVNG: p = 0.048; p = 1.00; p = 0.403, last 3 days respectively; Figure 6B). Neither group showed robust responding at the inactive nosepoke, and there were no significant differences between groups on inactive responses across the last three days of training (Day: F_2,36_ = 2.95, p = 0.065; Day x Treatment F_2,36_ < 1; Treatment F_1,18_ =1.969, p = 0.178). Control and EE+LVNG-treated animals did not differ when we assessed break point of the progressive ratio task for a grain pellet reward (t_17_ = 0.80, p = 0.43; Figure 6C).

**Figure 6.**
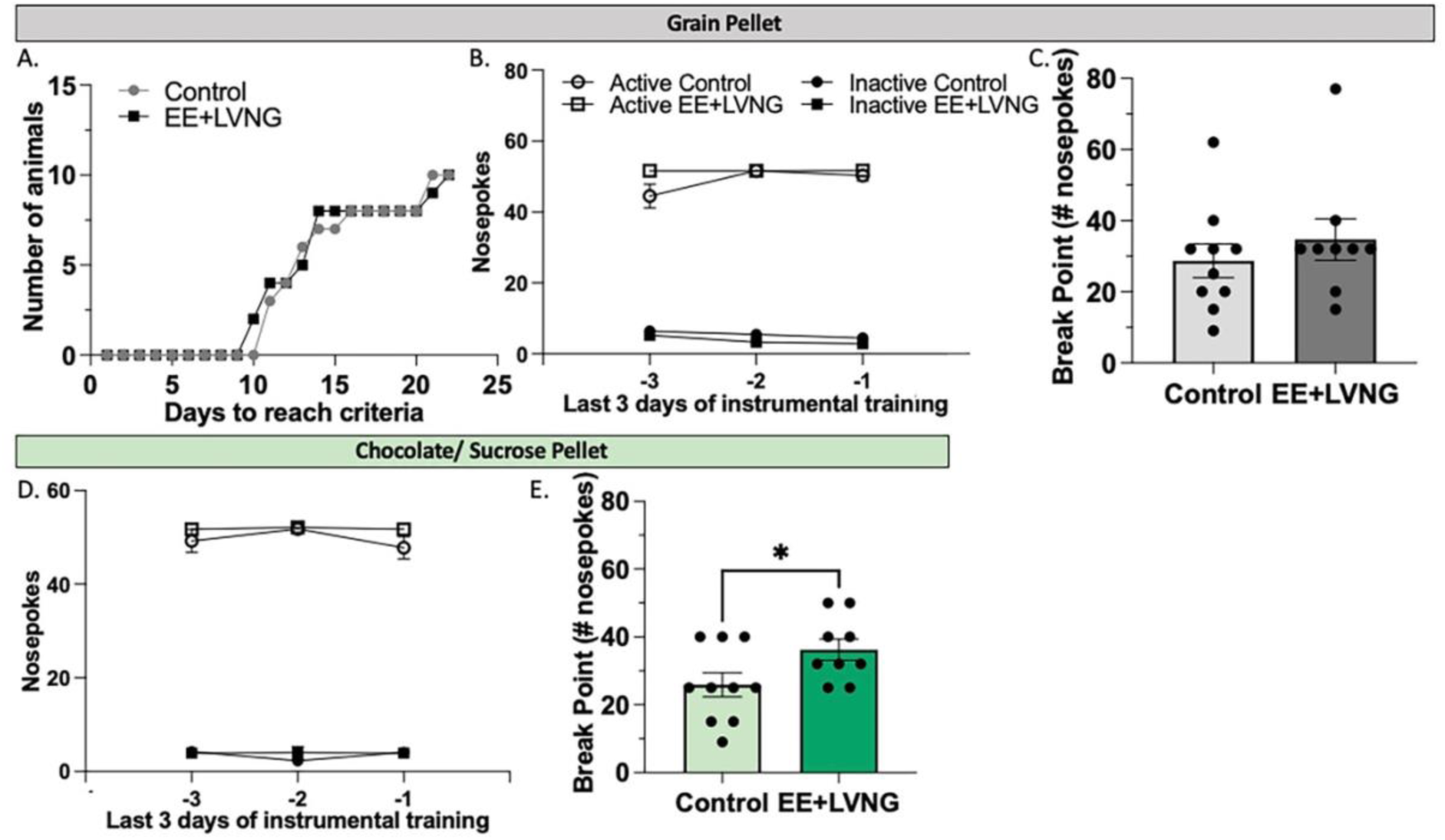
The sucrose preference findings could not be explained by a taste aversion nor a global decrease in motivation. (A) There was no difference in the number of days to reach criteria in instrumental training between treatment groups. (B) EE+LVNG-treated mice did not differ from control mice in number of active nor inactive nosepokes during the three instrumental training sessions prior to the progressive ratio with grain pellet rewards or (D) sucrose pellet rewards. (C) The EE+LVNG-treated animals did not differ from control animals in the progressive ratio task with grain pellet rewards; however, (E) EE+LVNG-treated animals had a higher break point compared to control animals in the PR task with sucrose pellet rewards, *p <0.05. Cohort 3: control n=10, EE+LVNG n=10. Light bars: Control-treated mice, Dark bars: EE+LVNG-treated mice.

When animals were retrained with sucrose pellets, there was no difference over the last 3 days of performing to criterion to the active nosepokes (Day F_2,35.95_ = 1.608, p = 0.21, Treatment: F_1_ = 1.023, p = 0.33; Day x Treatment: F_2,35.95_ = 1.158, p = 0.33) or inactive nosepokes (Day: F_1.84, 33.04_ = 0.997, p = 0.37, Treatment: F_1_ = 1.667, p = 0.73; Day x Treatment: F_1.84, 33.04_ = 1.912, p = 0.17; Figure 6D). Surprisingly, EE+LVNG-treated animals had a significantly higher break point than control animals with a sucrose pellet reward (t_17_ = 2.16, p = 0.045, η_p^2^_= 0.22; Figure 6E).

Together these findings demonstrate that EE+LVNG did not alter the reward-related learning processes in instrumental conditioning, nor did it affect motivation for food pellets, but EE+LVNG treatment did increase motivation to respond for sucrose. Thus EE+LVNG disruption of sucrose preference is not due to a global decrease in motivation.

## DISCUSSION

In this study, we characterize a new mouse model of OC exposure with translationally relevant dosing, designed to mimic daily oral contraceptive exposure in humans. Chronic daily EE+LVNG was well tolerated in mice, readily and rapidly ingested when given in 10% sucrose, and did not cause changes in weight gain. We observed suppressed estrous cycles in mice in this model, as assessed by vaginal cytology. EE+LVNG exposure caused a blunting of the corticosterone response to acute stress in EE+LVNG-treated animals without changes in basal corticosterone levels. In contrast, EE+DRSP-treated mice showed a robust and intact corticosterone response, suggesting that the blunted corticosterone response is not due to EE effects, but is rather specific to some classes of progestin. This observation is consistent with analyses of human data (Herrera et al., 2019) showing that LVNG-containing OCs show a stronger suppressive effect than other formulations. Behaviorally, we observed that chronic daily EE+LVNG caused a reliable, replicable, and specific anhedonia-like effect, without broad changes in anxiety, motivation, or depression-like behaviors.

How EE+LVNG and other OCs cause suppression of the acute stress response is a complex question. Here, corticosterone measurements – like cortisol measures in human studies – were taken at a single snapshot in time. Thus, lower levels might represent a suppressed HPA-axis response or may represent broader dysregulation of the HPA-axis. Nevertheless, our ability to mimic cortisol suppression in this mouse model provides a basis to further explore OC regulation of the HPA-axis from corticotropin releasing hormone in the paraventricular nucleus of the hypothalamus, to ACTH release from the pituitary, to peripheral corticosterone dynamics (Bangasser & Valentino, 2014; Heck & Handa, 2019). In particular, identifying downstream effects of stress on brain regions, including nucleus accumbens (Heshmati & Russo, 2015) and hippocampus (Wiborg, 2013), and molecular pathways including FKBP5 (Hartmann et al., 2021; Hertel et al., 2017) will be particularly relevant for understanding how OCs modify risks for anhedonia and depression. This model provides a platform to conduct mechanistic studies to assess OC’s effects on mood, behavior, and the brain.

In this mouse model, we observed specific anhedonia-like behavioral changes. In contrast, studies in rats (Follesa et al., 2002; Porcu et al., 2012; Simone et al., 2015) that have observed broader depression-like, stress-coping, and anxiety behaviors. This discrepancy is likely due to differences in doses, timing, and routes of administration (Tronson & Schuh, 2022). In this project, we intentionally used lower doses in mice than is typically used in rat models (e.g. Lacasse et al., 2022; Porcu et al., 2012) and we timed OC-administration to occur prior to lights out, and all testing was conducted at least 16 hours later. Further, in contrast with previous studies that have focused on persistent changes as a consequence of hormone exposure (Boi et al., 2022; Santoru et al., 2014), in our study, OC administration continued throughout the experimental timeline, including behavioral testing and stress. Given the complex effects of OCs, specific administration protocols, doses, and timing are likely to drive behavioral changes *via* different aspects of OC effects (Tronson & Schuh, 2022). For example, OCs exert effects *via* direct agonistic effects on estrogen and progesterone receptors (Fleischman et al., 2010; Graham & Milad, 2013; Porcu et al., 2012, 2019; Simone et al., 2015); and *via* suppression of circulating gonadal hormones including testosterone, progesterone, and estradiol that might persist after hormone exposure (Fleischman et al., 2010; Graham et al., 2018; Graham & Milad, 2013; Porcu et al., 2019; Pritschet et al., 2020). OCs also have indirect effects on androgenic, glucocorticoid, and mineralocorticoid receptors (Africander et al., 2011; Fedotcheva, 2021; Fuhrmann et al., 1996; Hertel et al., 2017; Kirschbaum et al., 1995; Phillips et al., 1990; Sitruk-Ware & Nath, 2010). Comparing across different rodent models with different administration protocols is therefore a critical advantage of animal work, as it allows us to disentangle different mechanisms of OC effects on stress-responses, brain, and behavior.

Our findings are consistent with those from human OC users – despite consistent suppression of the cortisol response to acute stress (Hertel et al., 2017; Kirschbaum et al., 1995; Merz & Wolf, 2017), most individuals do not experience major changes in mood, depression, or anxiety; nor does “the pill” cause major weight gain, despite persistent misconceptions (Brynhildsen, 2014; Gallo et al., 2014). Considering these observations in human subjects, our data provide strong support for the utility of this model and suggests that using lower, more translationally relevant daily oral dosing of EE+LVNG may effectively model OC use in humans. This model builds upon a growing recent body of literature identifying important effects of hormonal contraceptives on the brain, with both mood-related benefits for many (Keyes et al., 2013) and increased risks for depression in a subset of individuals (Poromaa & Segebladh, 2012; Schaffir et al., 2016; Skovlund et al., 2016; Worly et al., 2018). It adds to extant rat models of hormonal contraceptive exposure (Follesa et al., 2002; Porcu et al., 2012; Simone et al., 2015), which together demonstrate the importance of exogenous hormone treatment on cognition (e.g., Lacasse et al., 2022), affective behaviors (e.g., Heinzmann et al., 2014; Planchez et al., 2019), and stress (Porcu et al., 2019).

There are two important limitations that need to be addressed for further characterization of this model. First, as detailed above, to identify specific and detailed changes in HPA-axis function and implications for depression risk. Second to determine changes in ovarian hormone levels in response to OC exposure. Indeed, our reliance on vaginal cytology as our main measure of cycle suppression is a limitation of this project – particularly in mice for whom estrous cycles are easily disrupted (Lenert et al., 2021). Assessment of circulating estradiol, progesterone, or LH levels in mice is complicated: repeated, daily blood draws to determine the cyclicity is stressful for mice (Teilmann et al., 2014), which has its own effects on suppression of LH surges in mouse estrous cycles (Wagenmaker & Moenter, 2017). More importantly, estradiol assessment is often not reliable in mouse blood-draw volumes (Haisenleder et al., 2011; Wagenmaker & Moenter, 2017). Finally, although suppression of estradiol is well cited as a mechanism of OC efficacy, it is not clear they are consistently suppressed with OC use (Pritschet et al., 2020). Nevertheless, identifying OC-dependent changes in gonadal hormone will be a critical step towards identifying the precise mechanisms of action by which OCs regulate HPA-axis, modify anhedonia-like responses, and contribute to risk or resilience to depression- and anxiety-like behaviors.

In this paper, we begin to characterize a new mouse model of OC exposure that mimics key aspects of human OC effects. Together with other contraceptive hormone exposure models in rats (Porcu et al., 2019), this model provides essential tools with which to begin to examine questions arising from human studies on OC use that are neither feasible nor ethical to consider in human participants (Tronson & Schuh, 2022). These questions include the identification of individual differences that predict beneficial or detrimental mood effects of specific OCs and how contraceptive hormones modulate the HPA axis and blunt the stress response. Thus, this model—together with complementary animal and human studies—will be critical for identifying mechanisms and specific risk factors for adverse effects of OCs on mood. In the future, this work will be essential for developing an optimized and personalized approach to OC use to improve precision medicine.

## ACKNOWLEDGEMENTS

This work was funded by the Oscar Stern Award from the Eisenberg Family Depression Center at University of Michigan to NCT. Thanks to Jennifer Murray for edits and comments and undergraduate researchers Leah Conrad, Hannah Smith, Katie Alltop, Ikponmwosa Pat-Osagie, and Amy Choi for discussions on this project and assistance with behavioral tasks.

